# Evolution of burrowing and associated behavioral traits in Lagomorphs

**DOI:** 10.1101/2025.10.25.684593

**Authors:** Harsha Kumar, R Nandini

**Affiliations:** Indian Institute of Science Education and Research (IISER) Tirupati, Tirupati, Andhra Pradesh 517507, India

## Abstract

Burrowing is a complex behavior that has evolved repeatedly across animal lineages, yet its evolutionary origins and ecological correlates remain poorly understood in many mammalian groups. The order *Lagomorpha*, comprising pikas (*Ochotonidae*), rabbits and hares (*Leporidae*) exhibits remarkable diversity in lifestyle, sociality, and life history, making it an ideal system to study the evolution of burrowing behavior. Here, we reconstruct the most comprehensive dated phylogeny of Lagomorphs to date, encompassing 84 of 102 extant species, including newly sequenced genomes of *Ochotona nubrica* and *O. ladacensis*. Using ancestral state reconstruction and phylogenetically informed comparative analyses, we examine how burrowing behavior relates to morphology, climate, habitat, sociality, and fecundity. Our results reveal that burrowing evolved multiple times independently within Lagomorphs, with the ancestral state of Ochotonids inferred as burrowing and that of Leporids as semi-fossorial or form-dwelling. Contrary to expectations, burrowing species did not show distinct hind-limb adaptations or associations with extreme or variable climates. However, burrowing behavior was significantly correlated with sociality, and both burrowing and social species exhibited higher fecundity, suggesting an energetic and reproductive advantage linked to group-living in fossorial taxa. These findings highlight a recurrent evolutionary association between burrowing, sociality, and reproductive investment: the burrowing, sociality and fecundity triangle. This underscores the importance of integrating behavioral and ecological data to understand the adaptive evolution of lifestyle traits in mammals.

## Introduction

The animal kingdom has diverse examples of stunning architecture, such as beaver dams, nests of various species of birds, spider webs, beehives, and complex burrow systems of rodents (Gilliard 1963; Šklíba et al. 2012; Šumbera et al. 2012; Laidre 2021; Yang et al. 2022). These extended phenotypes or constructions are behavioral responses to environments that have evolved through natural selection and are predominantly innate behaviors (representative of a species/population) (Weber and Hoekstra 2009; DiRienzo and Dornhaus 2017). Such extended phenotypes can directly or indirectly affect the fitness (at the level of life history traits or through better protection from predators, increasing foraging efficiency, etc.) of architects in the wild and further drive their own evolution (Møller 1980; Pinter-Wollman 2015; Medina et al. 2022; Perez et al. 2020; Sugasawa and Pritchard 2022). A large-scale comparative phylogenetic study of such extended phenotypes of closely related species has the potential to address how such behaviors have evolved, their adaptive potential, their associations with life history/other behaviors, and potentially even uncover genetic elements that underline such evolutionary responses (Weber and Hoekstra 2009; Weber et al. 2013; Hu and Hoekstra 2017; Metz et al. 2017).

Members of the order Lagomorpha (*Leporidae*: Rabbits, Hares, and *Ochotonidae*: Pikas) are distributed across all continents except Antarctica, inhabit a wide range of habitats, have large variations in morphology (body size and appearance), social behavior (social, living in pairs, asocial), life history (species spread across the r-K spectrum of strategies) and lifestyle (Burrowing, Rock dwelling, forms, burrows of other animals, tree-holes, etc.,) (Alves et al. 2007; Smith et al. 2018). Lagomorpha is, therefore, a sound model system to study the evolution and adaptive significance of lifestyle and life-history traits given their diversity and low phylogenetic signal (similar traits found in distantly related species).

A small fraction of Lagomorphs are known to lead a semi-fossorial life. While some members of *Ochotonidae* construct both residential and breeding burrows using scratch-digging, others live and breed in rock crevices (Alves et al. 2007; Smith et al. 2010; Wilson and Mittermeier 2016; Smith et al. 2018). A large fraction of species from the family *Leporidae* either construct forms (shallow scrapes in soil that appear cup-like over repeated use)/ use burrows of other mammals for residence but construct rudimentary burrows/nests above ground with vegetation and fur while breeding (Wilson and Mittermeier 2016; Smith et al. 2018). A small fraction of species in both *Leporidae* and *Ochotonidae* are known to have flexible lifestyles allowing them to utilize/ construct multiple refuge types for residing and breeding (for instance: *Ochotona rufescens* is known to live and breed in both rock crevices but also construct burrows on steppes; *Sylvilagus audubonii* is known to construct nests above ground while breeding; but construct forms or uses burrows of other species on extremely hot days) (Smith et al. 2018; Fulk and Khokhar 1980; Schmidly and Bradley 2016).

Burrowing species are known to have morphological adaptations in tibial (associated with scratch-diggers), mandibular morphology (associated with chisel-tooth diggers) and cranial morphology to facilitate fossorial life, as shown in many rodents (Gomes Rodrigues et al. 2016; Das et al. 2022; Gomes Rodrigues and Damette 2023; Gomes Rodrigues et al. 2023) and beetles (Emlen and Keith Philips 2006; Macagno et al. 2016). Species pre-disposed to burrowing behavior in Lagomorphs are expected to have large hind legs in proportion to their body, given that they are scratch-diggers (Hypothesis 1).

Burrowing is typically an energetically expensive behavior that has evolved multiple times in the animal kingdom (Kinlaw 1999), including many rodent model systems such as pocket gophers (*Thomomys bottae*), common degu (Ochoton degus), desert golden mole (*Eremitalpa granti*) and Mole rats (*Cryptomys damarensis, Heterocephalus glaber*) with its costs being particularly large in arid soils with low moisture (Vleck 1979; Lovegrove 1989; Ebensperger 1998; Seymour et al. 1998; Ebensperger and Bozinovic 2000). Group living is thought to facilitate the reduction of costs associated with burrowing in rodents (Spinks 1998; Ebensperger and Bozinovic 2000; Ebensperger and Cofré 2001; Lacey and Wieczorek 2003), wombats (Walker et al. 2007) and lizards (McAlpin et al. 2011). Sociality is, therefore, likely to be associated with burrowing behavior, particularly in arid landscapes (Hypothesis 2).

Thermoregulation can dominate the energy budgets of even small endothermic mammals living in highly seasonal environments or extreme environmental regimes when there are constraints on energy acquisition and expenditure (Speakman 1997; Milling et al. 2018). Extended phenotypes, such as burrows, not only provide stable thermal refuges to animals by buffering fluctuations in ambient temperature but also help animals meet their energetic requirements behaviorally without requiring a change in physiology (Pike and Mitchell 2013; Millar et al. 2016; Milling et al. 2018). Many species of temperate mammals, such as the steppe marmot (*Marmota bobax*) and the speckled ground squirrel (*Spermophilus suslicus*), are known to construct deep burrows that remain above-zero temperatures through the winter, allowing them to conserve energy during hibernation (Nikol’skii and Khutorskoi 2001; Belovezhets and Nikol’skii 2012). Such extended phenotypes are considered to be under selection, and we hypothesize that they are likely to be associated with extreme temperature regimes (hot or cold; Hypothesis 3.1) or highly variable seasonal temperature regimes (Hypothesis 3.2).

Predation risk is known to alter the behaviors of prey animals and can lead to varied responses, with some species/individuals avoiding predators altogether (change in activity patterns), others becoming vigilant and fleeing upon detecting the predator or concealing themselves through in a multi-stage response (Lima and Dill 1990; Suselbeek et al. 2014). Refuges such as burrows, tree hollows, and vegetation cover help animals avoid predators while continuing to forage with minimal daily risk (Hemmi 2005a; Hemmi 2005b; Wilson et al. 2012). Fidler crabs (*Uca vomeris*) and woodchucks (*Marmota monax*) that were closer to their burrows had short flight initiation distances and their responses were additionally modulated by the approach velocity of predators (Bonenfant and Kramer 1996; Hemmi 2005a; Hemmi 2005b). Voles (*Microtus spp*) placed in predator-free pens had fewer burrow entrances and fewer simple burrow systems when compared to pens that predators could access (Harper and Batzli 1996). It is expected that burrowing behavior is associated with largely open habitats if it is associated with predator avoidance (Hypothesis 4; Metz et al. 2017; Jackson 2000).

Life history theory tries to explain the diversity of reproductive strategies by placing emphasis on how ecology/environment shapes animals to partition limited energy acquired for survival and reproduction to optimize fitness (Stearns 1977; Fabian 2012). Life history traits are also subject to allometric scaling size, with small body-sized animals being more fecund and short-lived at the expense of parental care (r selected) (Pianka 1970; Blueweiss et al. 1978; Sibly and Brown 2007) (Hypothesis 5.1).

Climatic envelopes are known to control the number and timing of reproductive investments of species across many taxa (Post et al. 2001; Evans et al. 2005; Pankhurst and Munday 2011; Wells et al. 2022). In particular, extreme environments (hot/cold/variable temperature regimes and very wet or very arid landscapes) can constrain the expression of life history traits by allowing for relatively few breeding efforts during the year but with large investments in one bout of reproduction (Evans et al. 2005; Hufnagl et al. 2011; Battistella et al. 2018; Verhagen et al. 2020) (Hypothesis 5.2).

Sociality is known to affect the expression of life history traits in ground squirrels and canids, with social species being large body sized, associated with habitats having shorter growing periods and exhibiting delayed dispersal, reproduction although with larger litter sizes and survival (Armitage 1981; Bekoff et al. 1981). Although studies show varied fecundity responses due to sociality (ranging from positive to negative), it is now accepted that sociality has modest positive effects on fecundity (Ebensperger et al. 2012) (Hypothesis 5.3).

General observations of two well-studied pika species (*O. curzoniae*: burrower and *O. princeps*: rock-dweller) have led to the hypothesis that rock-dwelling species may be asocial in addition to being k-selected (long-lived, low fecundity, late maturity) in comparison to burrowing species which are social in addition to being r-selected (short-lived, high fecundity and early maturity) (Alves et al. 2007) (Hypothesis 5.4).

We build the first comprehensive dated phylogeny of Lagomorpha covering 81/109 extant species distributed across the globe by combining previously published exome data for *Ochotonidae* and multi-locus markers for *Leporidae* in this study. In the process, we also solve the disputed phylogenetic placement of *Ochotona ladacensis*.

We collected natural history data for these species of Lagomorphs using field guides, research articles, museum databases, and government portals to address the hypothesis outlined above. We use the comparative phylogenetics method in conjunction with ancestral state reconstructions to understand how burrowing behavior has evolved in Lagomorphs. We explore if burrowing in contemporary timescales is associated with sociality (burrowing is energetically expensive) (Hypothesis 2), extreme/variable climatic regimes (Hypothesis 3.1, Hypothesis 3.2), open habitats, and diurnal activity periods (Hypothesis 4.1, Hypothesis 4.2) and hind-limb morphology (scratch-diggers) (Hypothesis 1). Furthermore, we examine if burrowing species are more fecund (Hypothesis 5.4), social species are more fecund (Hypothesis 5.3), and small body-sized animals are more r-selected (allometric scaling; Hypothesis 5.1). Additionally, we explore if climate (temperature and precipitation) can directly drive fecundity patterns of Lagomorphs across the world (Hypothesis 5.2). This study addresses the evolution of lifestyle and fecundity patterns of Lagomorphs and the ecological factors that drive their expression.

## Methods

### Whole genome sequencing of Ochotona nubrica and Ochotona ladacensis

We obtained one tissue sample of each species while trapping and ear tagging animals on select species colonies at the Changthang Biotic province, Ladakh, India.

These tissue samples were stored in absolute alcohol at -20°C until further processing. DNA was extracted from these tissues using a Qiagen Blood and Tissue Kit following a modified protocol. These changes from the standard protocol included the addition of RNAase and the introduction of a second digestion step at 70°C with the addition of AL buffer. Column purification and Illumina Sequencing at 30x (∼2GB genomes) was outsourced to Neuberg Supratech Reference Laboratory and resulted in the generation of ∼60GB of data per species. The raw reads obtained from the sequencing were processed in the ways described below to place these species on a phylogeny.

### Tip Data collection

Natural history data available on lifestyle (residential and breeding habits), habitat preferences, activity patterns, and social behavior was collected and compiled from credible published literature, such as books, field guides, several research papers, and museum websites, for species of the order *Lagomorpha* (89/102 extant species). Life history data (number of offspring/litter, number of litters/year, total number of offspring/year) and morphometric data (body size, body mass, and hind-foot length) were also collected to test the hypotheses enumerated. Each species’ length of the breeding season and gestation period were collected to assess the validity of the fecundity data gathered from field guides and books. This is the first dataset of lifestyle, life history, and other morphometric traits put together by carefully gathering and combing data across sources and validating them. Naturally, way too many references were involved in creating this database, and it is impossible to cite all references in the text here (provided in Supplementary Table 1).

To understand if ecology drives lifestyle responses of *Lagomorphs*, we downloaded all open source CHELSA bio-climatic data for the time frame (temperature and rainfall-related variables for the time frame between the years 1973 and 2013: Bio1- Bio19) available at a spatial resolution of 1kmsq and extracted data from grids that overlapped shape layers obtained from IUCN. After cleaning the climatic dataset to remove empty cells (limitations of satellite-acquired data), each species’ mean, minimum, and maximum values for each bio-climatic variable were calculated in its distribution range. Furthermore, introduced populations of *Leporidae* species were ignored, and only native ranges were considered while computing climatic envelopes of species.

A pseudo-continuous metric that captures the energetic cost of burrow construction and levels of pre-disposure to the activity (Burrowing predisposition) was constructed following published work (Noonan et al. 2015). On this metric, primary burrowers (construction and burrow maintenance) were scored the highest: 1, secondary modifiers: 0.75 (augmenting and maintaining burrow systems), tertiary occupants (use pre-existing burrows): 0.5, forms (a most rudimentary form of digging): 0.25, life in tree holes and cavities: 0.125 and life in rocks : 0. Furthermore, since the breeding and residential lifestyles of many *Leporidae* members are different, the metric was evaluated separately for both breeding and residential lifestyles and used later for regressions against different ecological factors outlined in the hypotheses.

A second metric that captures habitat openness was also collated by collecting habitat information from various sources (Supplementary Table 1) for all species and ranking them based on sunlight penetration (Alpine meadows [7] > Grassland [6] > Rocks in open [5] > Alpine scrub [4] > Savannah [3] > Reeds [2.5] > Forest [2] > Rocks under forest canopy [1] > Forest Understory/Swamp Forest [0]). When a species occupied more than one habitat, an average of corresponding scores were taken to reflect states in between.

### Climatic data processing

Bio-climatic data (World Clim dataset: Bio1-Bio19 representing different temperature and precipitation variables) was collected at the spatial scale of the world at a 1km*1km resolution. Using the distribution shape files obtained from IUCN as a reference, data was extracted from the species’ distribution range using the Zonal Statistics plugin on QGIS to give pixel values corresponding to mean, median, maximum, and minimum.

It is to be noted that the data collected was further cleaned to exclude species with very large distribution ranges and varied chromosome numbers (left with 84/102). All Quantitative data was handled considering repetitive data (duplicates across literature sources were considered only once). All morphometric data (only the mid-point was considered for range data) was averaged, and the Z-score was transformed within the families *Leporidae* and *Ochotonidae* to account for the difference in body size across different Lagomorphs.

### Tree building

The raw reads for the exome dataset of the family Ochotonidae (NCBI accession numbers SRR10107529 - SRR10107633; Wang et al. 2020), Illumina data generated for two species of pikas (*Ochotona nubrica and Ochotona ladacensis*) and multi-locus markers for the family Leporidae (Melo-Ferreira et al. 2015) were downloaded and processed separately and stitched together to construct a well resolved phylogenetic tree of the order Lagomorpha. Additionally, genomes of two different species of pikas (*Ochotona ladacensis* and *Ochotona nubrica*) were sequenced to complete the phylogeny of pikas for ancestral state constructions of burrowing.

To build the *Ochotonidae* tree, raw reads were split (paired-end data) and downloaded using the fastq-dump of the SRA toolkit (version 2.10.2). All raw read data was subject to removing Illumina adapters and quality trimming from ends at a cut-off of phred score 30 (0.1% base-calling error rates) using Trim Galore (version 0.6.5). The American pika cds (version and accession on NCBI; Author) were downloaded to reference alignments of pika raw reads from Wang et al. 2020. A new, additional reference for the American pika was created manually to include multi-locus markers (ALB, DARC, OXA1L, PPOX, PRKCI, SPTBN1, TSHB, UCP2, and UCP4) available for species of pikas that weren’t on the exome dataset (*Ochotona illiensis, Ochotona rufescens, Ochotona rutila, Ochotona collaris, Ochotona argentata, Ochotona alpina, Ochotona hofmanni*). Most missing species were covered by pooling all NCBI sequences available across markers, with intronic and exonic regions being treated as separate reads to aid alignments (also done while editing reference cds). The multi-locus markers and trimmed raw reads (exome dataset) were aligned to their respective reference cds using bwa mem (Version 1.11) and followed by conversion of sam files to bam files. These bam files were sorted and indexed using samtools (Version 1.11), pooled across samples for a species using samtools mpileup and followed by variant calling using bcftools (Version 1.11) at a minimum depth cut off of 3 and maximum depth cut off of 100. Variant call files generated were sorted, indexed and merged using vcf-merge (GATK version 4.1.4.1). In addition, raw read data for *Ochotona princeps* (NCBI accession number: SRR409146) and *Mus musculus* (NCBI accession number: SRR327346) was also downloaded and processed as described above to introduce the tip used as reference and to root the tree, respectively. Quality control was ensured at each step using fastqc (version 0.12.1) before aligning reads to references, the integrative genomics viewer (version 2.16.2) post-generation of bam files, and manual inspections on Unix shell. The merged VCF file was converted to phylip format accessible by maximum likelihood tree builders such as iqtree (Nguyen et al. 2015; Chernomor et al. 2016; Kalyaanamoorthy et al. 2017; Hoang et al. 2018; Minh et al. 2020)(Version 2.0). An ultrafast bootstrap *Ochotonidae* tree (10000 runs with each site being modeled by a different evolutionary rate and sh-alrt bootstrap test: 100 runs) was generated with *Oryctolagus cunicularis* as an outgroup. This generated a species-level phylogeny that was well resolved and placed missing species on the phylogeny allowing for robust ancestral state reconstructions (node bootstaps > 95) (Figure 1).

**Figure 1.**
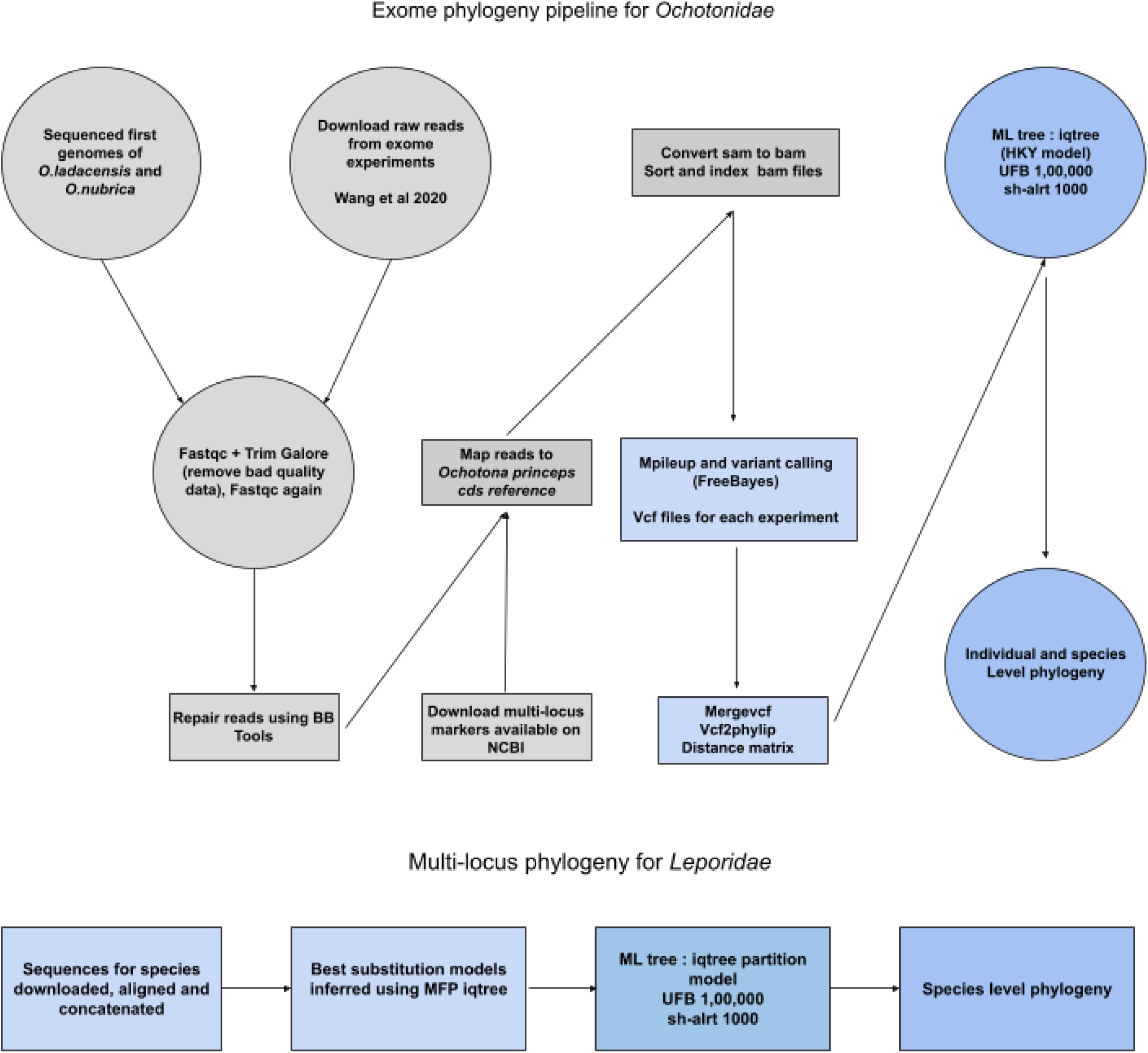
: Pipeline used to construct Lagomorpha dated phylogeny.

To construct *Leporidae* tree, multilocus markers were downloaded from NCBI and aligned using Geneious Prime (Version 2020.2.4). Alignments generated for each marker were concatenated. The concatenated alignment was used to construct an ultrafast bootstrap *Leporidae* tree (1,00,000 runs with each marker being modeled by a different evolutionary rate with sh-alrt 1000 for estimating bootstrap supports) with *Ochotona princeps* as the outgroup (Figure 1). The two trees generated were stitched together at the Ochotonidae-Leporidae split. Branch lengths were calibrated using published primary (fossil data) and secondary divergence (from another study) time estimates to account for disparate datasets (Rodentia and Lagmorpha divergence - 87.8mya, Lagomorpha divergence - 60mya, MRCA of extant Ochotonidae - 13.75mya, MRCA of extant Leporidae - 18.1mya, MRCA of clade Ochotona - 10.46mya, MRCA of clade Conothoa - 7.5mya) (Ge et al. 2013; Wang et al. 2020).

### Ancestral Reconstruction of Lifestyle and Morphometric Traits Lifestyle

Since residential and breeding lifestyles of many species are starkly different, both lifestyles were considered in the study and characterized by different combinations of refuge use and construction as described. For ancestral state reconstructions, the following trait states were considered:

a. Four different states based on maximum energy expenditure by each species (burrowing > burrows of others > forms > rocks). If a species is known to construct its own burrows and use burrows of other species and forms, it is evaluated as a burrowing species representing the most energetic behavior exhibited by the species. This allows for visualization of jump-in energy states of behavior along the phylogeny.
b. Eight different categories (Burrowing, Rocks, Forms in vegetation, Intermediate 1 - Rocks + Burrows of other animals, Intermediate 2 - Forms in vegetation + Burrowing, Intermediate 3 - Forms in vegetation + Rocks, Flexible 1 - Rocks + Burrowing, and Flexible 2 - Forms in vegetation + Burrows of other animals/own burrows + Rocks) representing the most descriptive account of all combinations of trait states.
c. Two categories (burrowing and non-burrowing) representing the most simple trait states. Here, only if an animal constructed its own burrows it was considered a burrowing species.

Different hierarchical versions of the Mk model (ER, SYM, and ARD) were run to select the best approximation of the evolution of lifestyle (Revell 2012). The best model selected was used for stochastic character mapping (insert citation) of lifestyle traits on the phylogeny. Transition rates between states were generated using the simmap function on the ape package (Paradis and Schliep 2018).

Morphometric traits were modeled as a Brownian motion process over phylogenies and mapped over branches to visualize body size and body mass evolution over large time scales.

### Phylogenetic Generalised least squares and Phylogenetic independent contrasts

We performed phylogenetically corrected regressions and phylogenetic independent contrasts with multiple predictor variables (bio-climatic variables, morphology, habitat) and one dependent variable (residential lifestyle, breeding lifestyle, and life history) using the comparative phylogenetics framework to understand drivers of lifestyle and life history at a macro-evolutionary scale. R studio (version 4.1) packages ‘caper’ (Orme 2013), ‘phytools’ (Revell 2012), ‘geiger’ (Pennell et al. 2014) and ‘ape’ (Paradis and Schliep 2018) were used to import trees, tip data and perform regression analysis.

## Results

### Lagomorpha phylogeny

We constructed a well-resolved dated maximum likelihood phylogeny of Lagomorpha (90% of nodes with bootstraps > 90), including more than 2/3rds of species from the order (Figure 2). We placed the population of *Ochotona nubrica* from the Changthang Biotic Province, Ladakh, India, basal to all other *Ochotona nubrica* in the clade *Ochotona*. The population of *Ochotona ladacensis* from the province was placed in the clade *Conothoa* basal to the *Ochotona himalayana, Ochotona macrotis,* and *Ochotona roylei* group.

**Figure 2:**
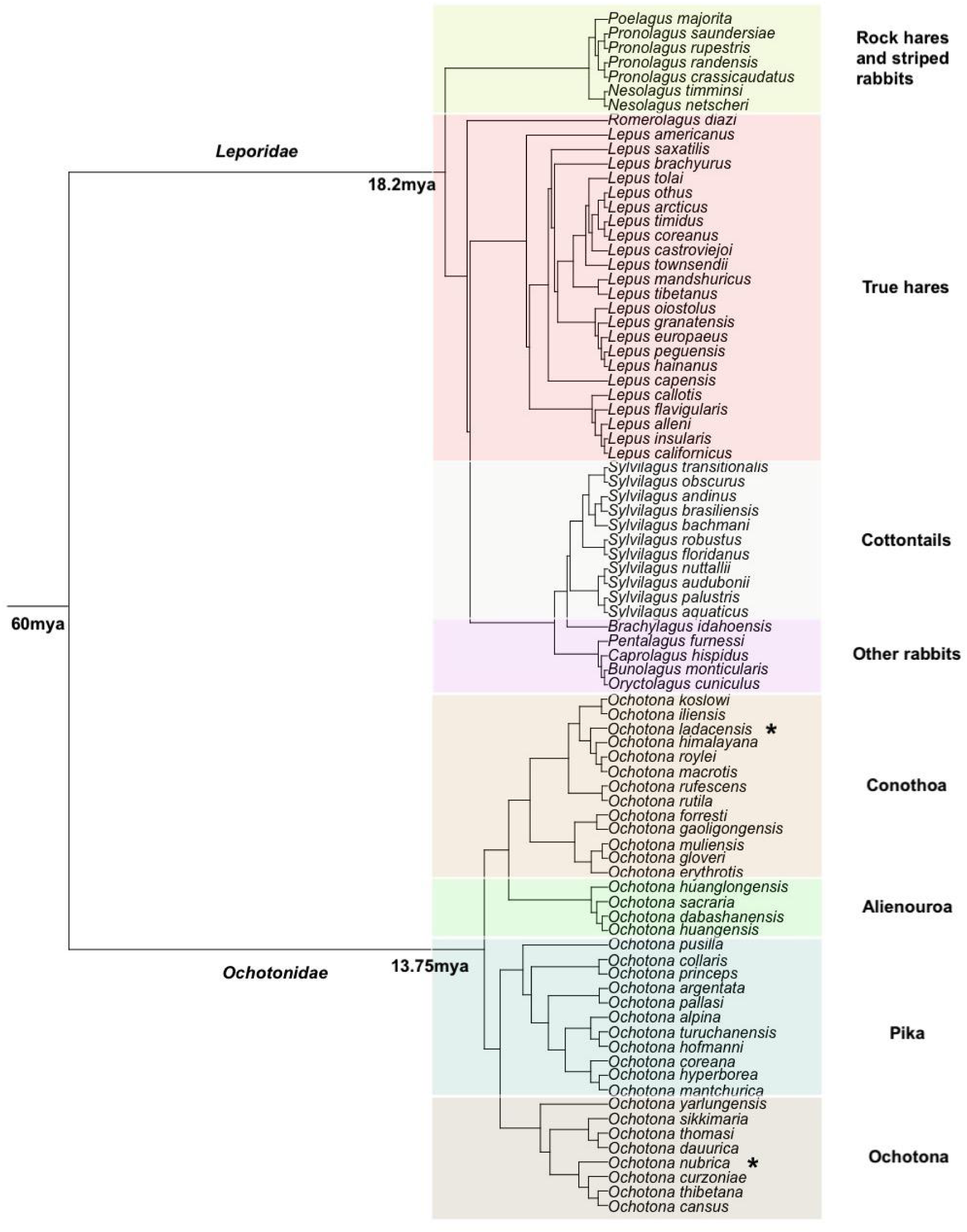
Well-resolved phylogeny of Lagomorpha (90% nodes with bootstraps > 90) constructed using a combination of exomes and multi-locus markers. Species annotated with stars represent species placed on phylogeny with whole genome sequencing in this study.

### Ancestral state reconstruction: Morphometric and Lifestyle traits

Lifestyle - The ancestral state of all *Ochotonids* was inferred to be in the burrowing state, while that of *Leporids* was either living in forms/burrows of other animals when four different residential lifestyle categories based on maximum energetic investment by species (burrowing > burrows of others > forms > rocks) were considered (Figure 3). The ancestral state of all Lagomorphs was not resolved. The ancestral states recovered were similar when two or eight residential lifestyle categories were considered for each species (see methods for treatment of lifestyle traits) (Figure S1, Figure S2).

**Figure 3:**
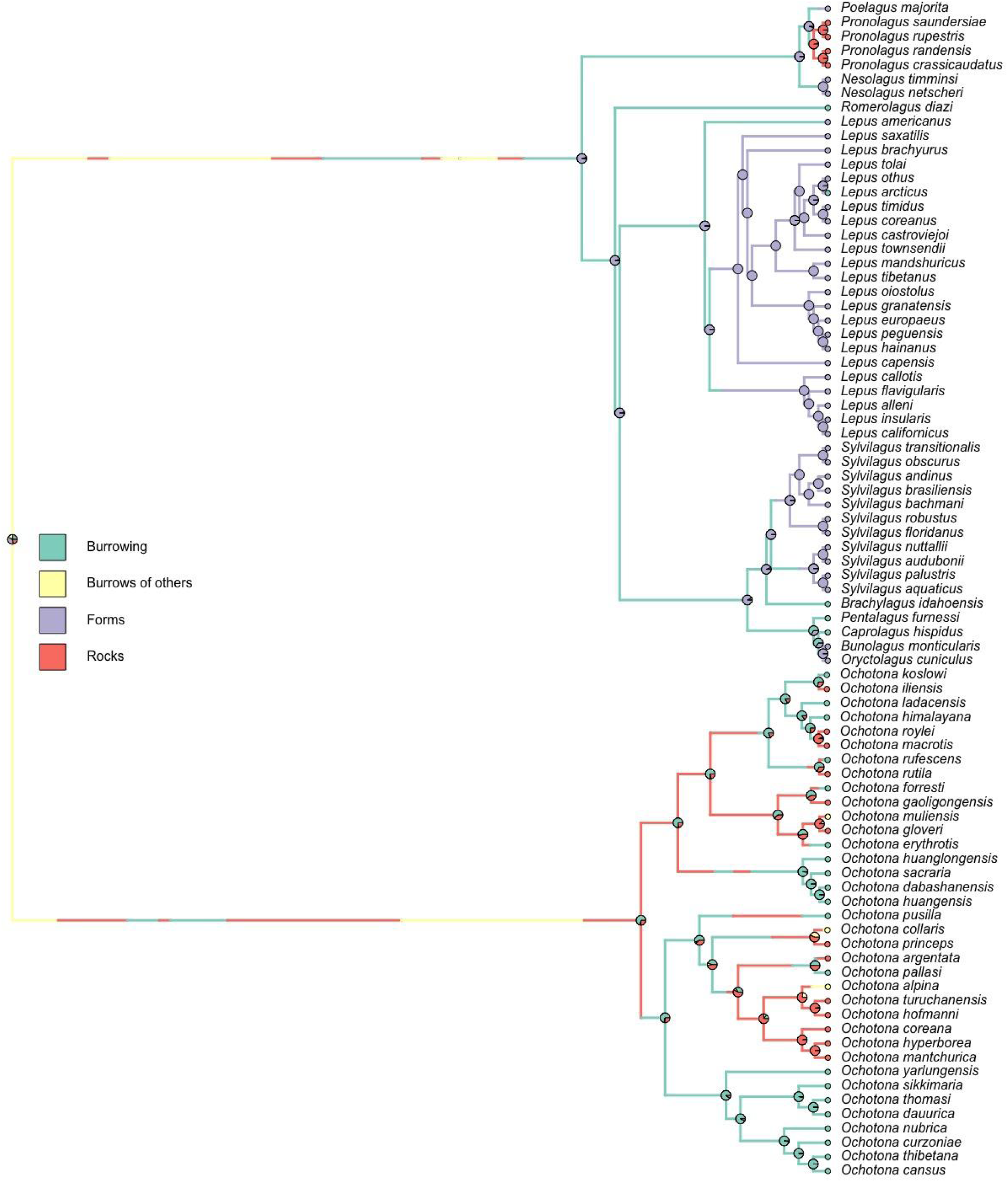
Ancestral state reconstruction of residential lifestyle on Lagmorpha phylogeny. The tips have been colored based on lifestyle traits, and pies at nodes are colored based on the probability of trait states based on stochastic character mapping.

A summary of stochastic character maps revealed that the most common transitions to burrowing behavior included those from living in rocks (26% of all transitions), as seen in the sub-clade *Conothoa* of *Ochotonidae* and living in forms (12% of all transitions) in *Leporidae* (Figure 4). Additionally, other common transitions included those from: a) burrowing to rock-dwelling behavior (29%) multiple times in *Ochotonidae* and once in Leporidae (rock hares and stripe rabbits); b) living in burrows of other animals to rock-dwelling (9%); c) rock dwelling to living in burrows of other animals (13%) (Figure 4).

**Figure 4:**
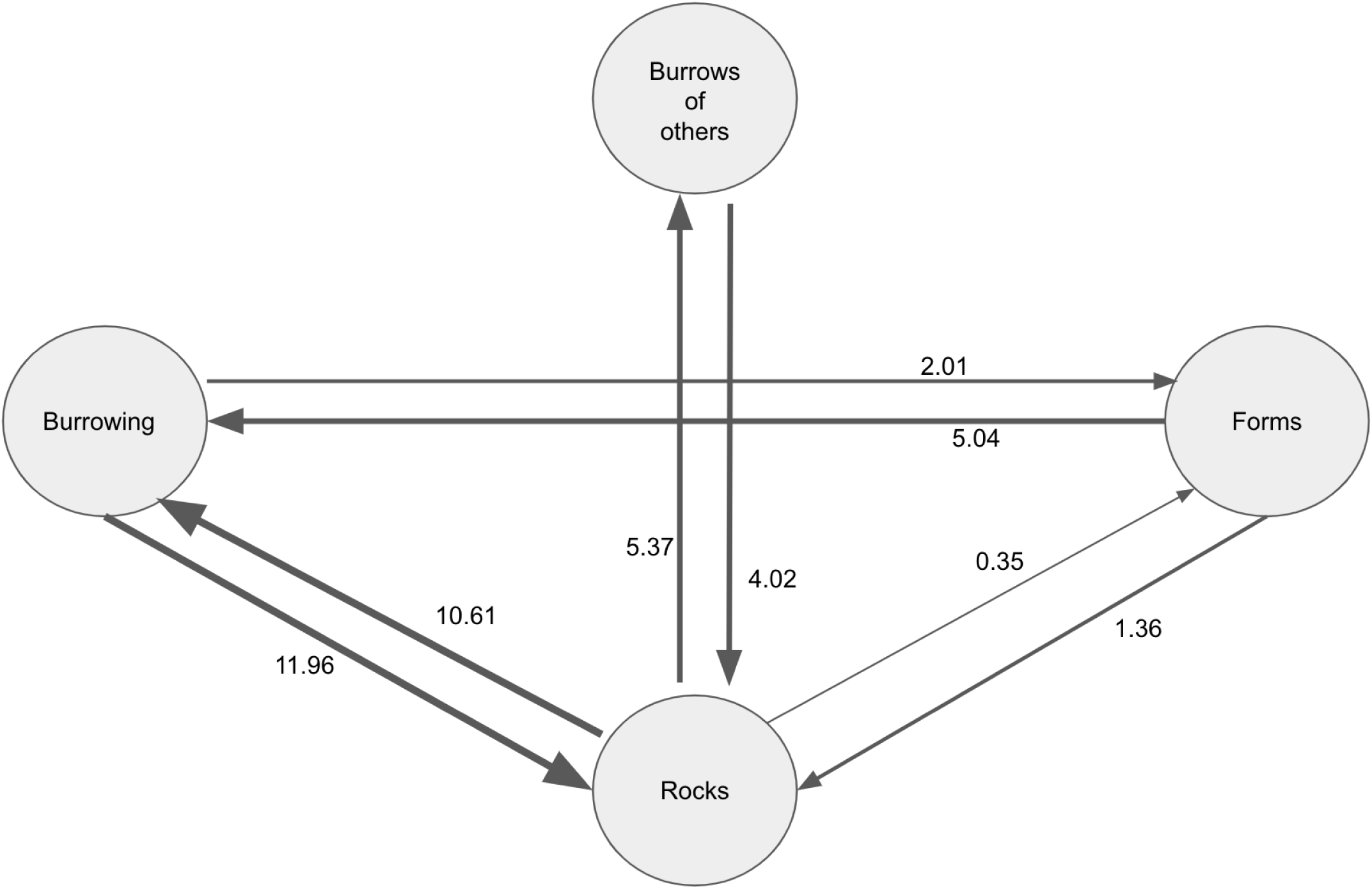
Transitions between lifestyle states as described by stochastic character maps. An average number of 40 transitions were identified across the phylogeny and the flowchart above documents the number of transitions between states (the thickness of the arrows depicts the percentage of all transitions).

#### Body size and Body mass

Ancestral state reconstructions indicated that the ancestor of all Lagomorphs had an intermediate body size. Subsequently, both small body size and large body size have evolved once in pikas and hares, respectively (Figure 5). Although ancestors of Lagomorphs had intermediate body mass, large species of Ochotonids are similar in body mass to small species of Leporids (Figure S3).

**Figure 5:**
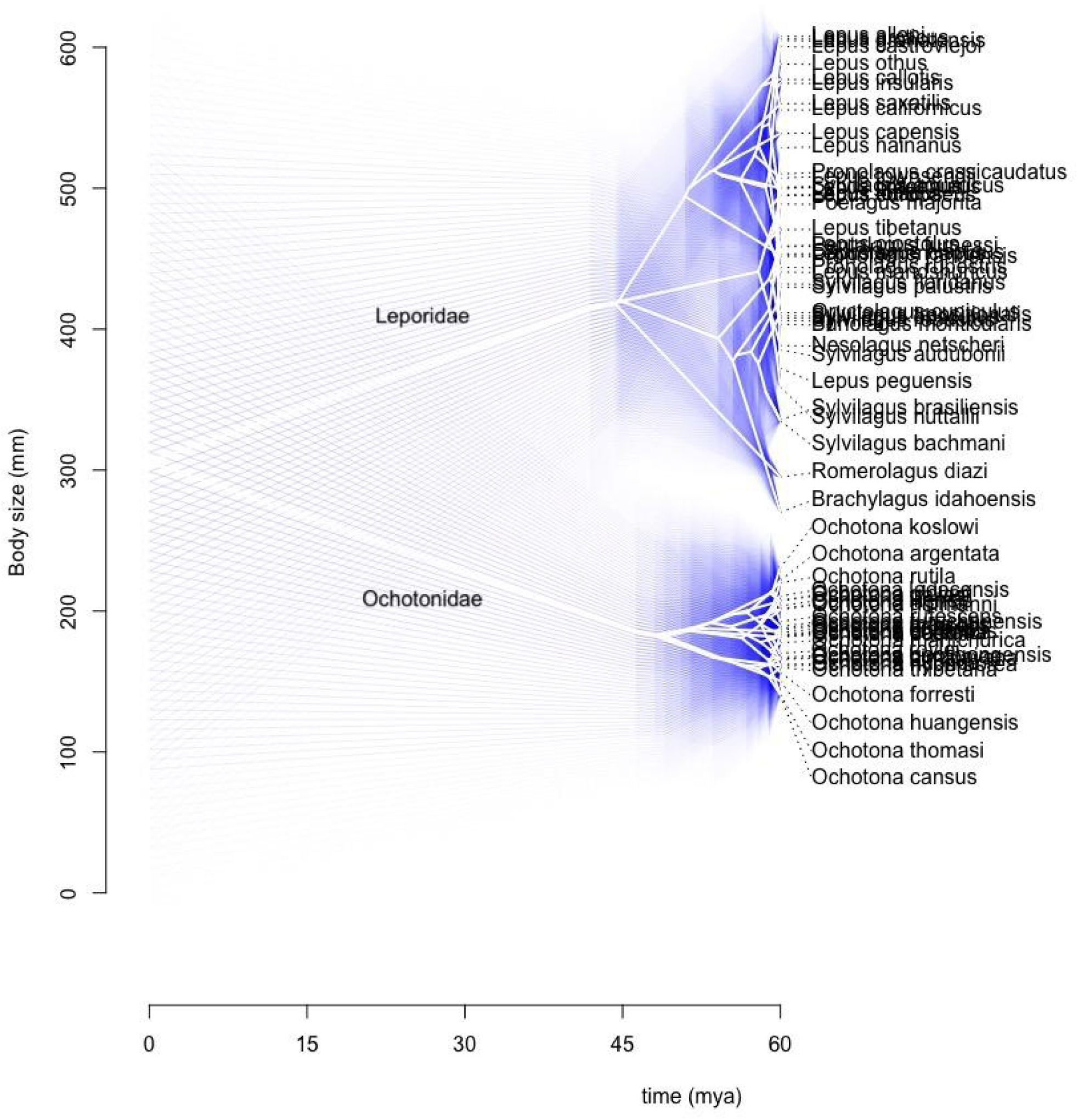
Phenogram of body size changes along phylogeny with 95% confidence intervals overlaid. *Ochotonidae* - Small body-sized, *Leporidae* - Intermediate body-sized: Rabbits, Large body-sized: *Lepus* hares.

#### Sociality

The ancestral state reconstructions for social behavior were not resolved due to the dispersed appearance of the trait along the phylogeny (Figure S4).

### Associations of lifestyle with morphology, sociality, climate, and predator avoidance

Lifestyle and morphological characteristics (Hypothesis 1) - Species of burrowing Lagomorphs did not have larger hindlegs relative to their body size as expected (Figure 6). A phylogenetic ANOVA revealed no significant differences in relative hindlegs between groups of Lagomorphs engaging in different lifestyles (burrowing, burrows of others, forms, rocks) (F-value: 16.22, p-value: 0.19682).

**Figure 6:**
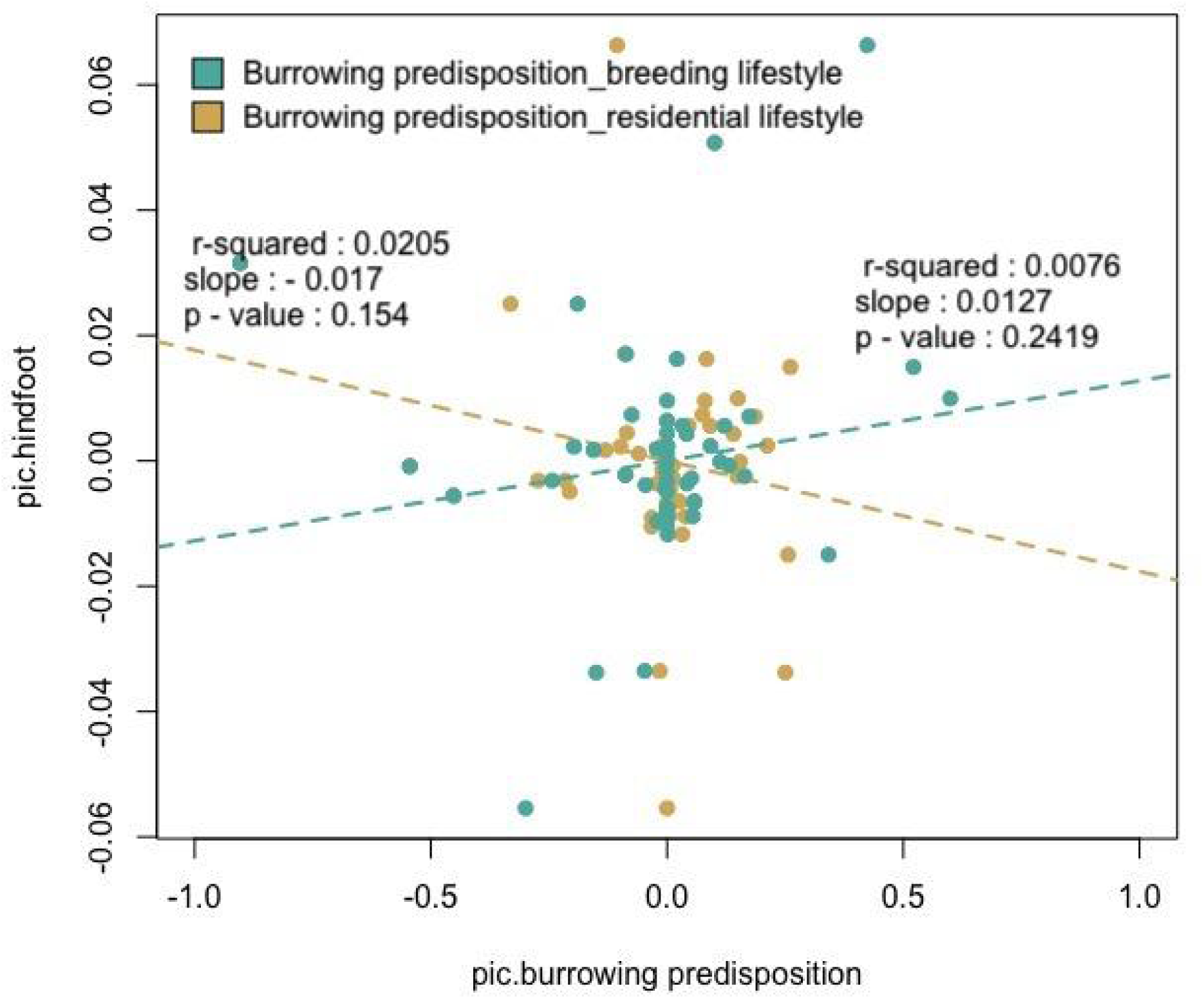
Phylogenetically controlled regressions of relative hindfoot length against burrowing predisposition.

#### Lifestyle and Sociality (Hypothesis 2)

Burrowing Lagomorphs were also social irrespective of whether residential or breeding lifestyles were considered (Figure 7; PIC_Burrowing propensity_Residential lifestyle vs. PIC_Sociality- Slope: 1.02, r-squared: 0.22, p-value < 0.05; PIC_Burrowing propensity_Breeding lifestyle vs. PIC_Sociality- Slope: 0.865, r-squared: 0.19, p-value < 0.05). This reconciles well with species’ natural history collated with all pikas and true rabbits that construct burrows being social, social hares (small group sizes) showing varied lifestyles, such as constructing burrows in extreme weather (*Lepus arcticus*), living in burrows of other animals (*Nesolagus netscheri*), in forms (*Sylvilagus aquaticus*) or even rocks (*Pronolagus randensis*) (Table S1_Raw Data).

**Figure 7:**
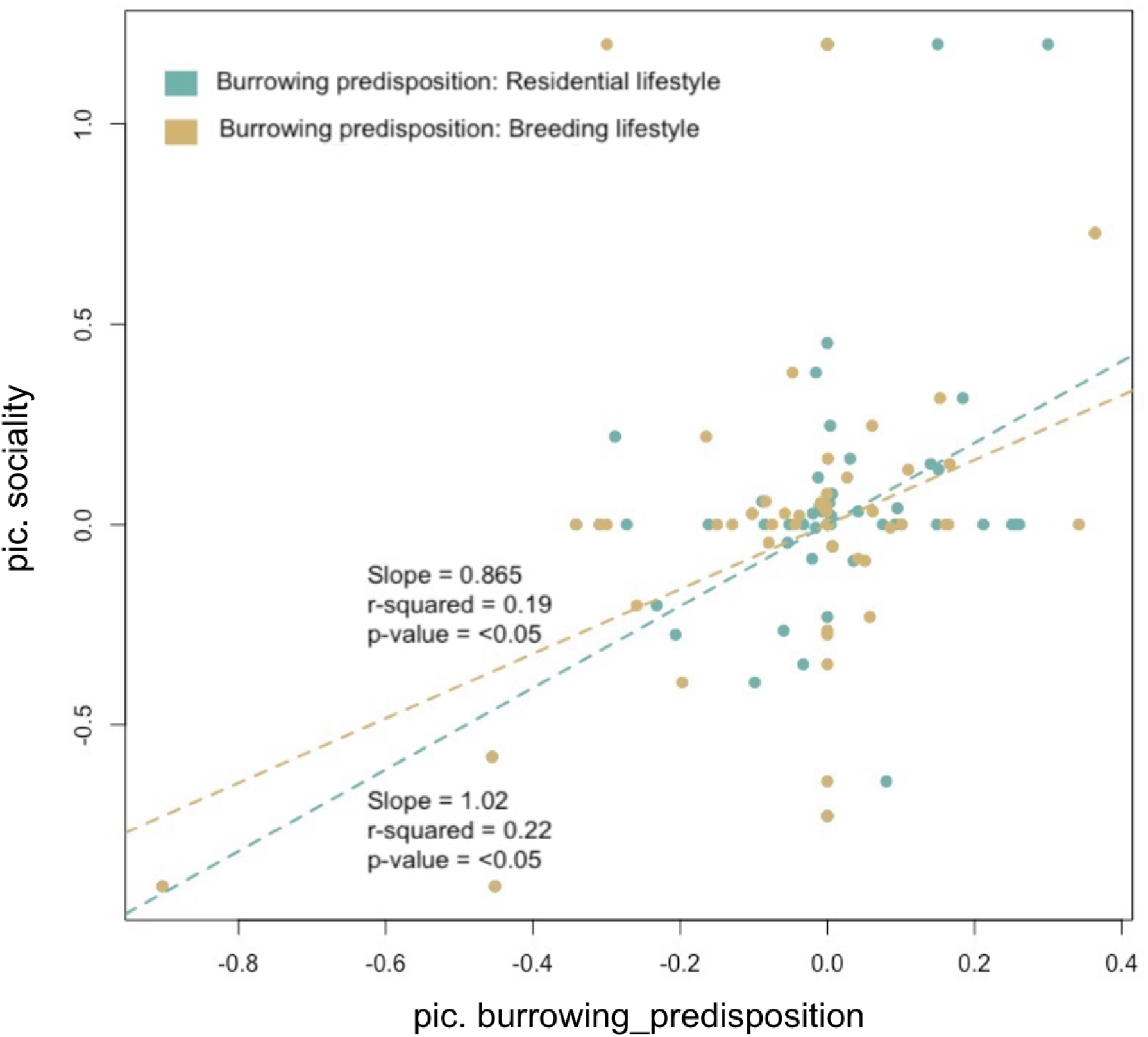
Phylogenetically controlled regressions of burrowing predisposition against sociality.

#### Lifestyle and climate (Hypothesis 3.1, 3.2)

Burrowing Lagomorphs were not associated with more extreme/ seasonal climates than other species with different residential lifestyles (Figure 8). All relationships were characterized by low r-squared, low slope, and insignificant p-values. When four lifestyle states were considered, no differences were found between groups for climatic variables even after controlling for phylogeny (Phylogenetic ANOVAs characterized by F-values: 0.6 - 12, and p-value > 0.4).

**Figure 8:**
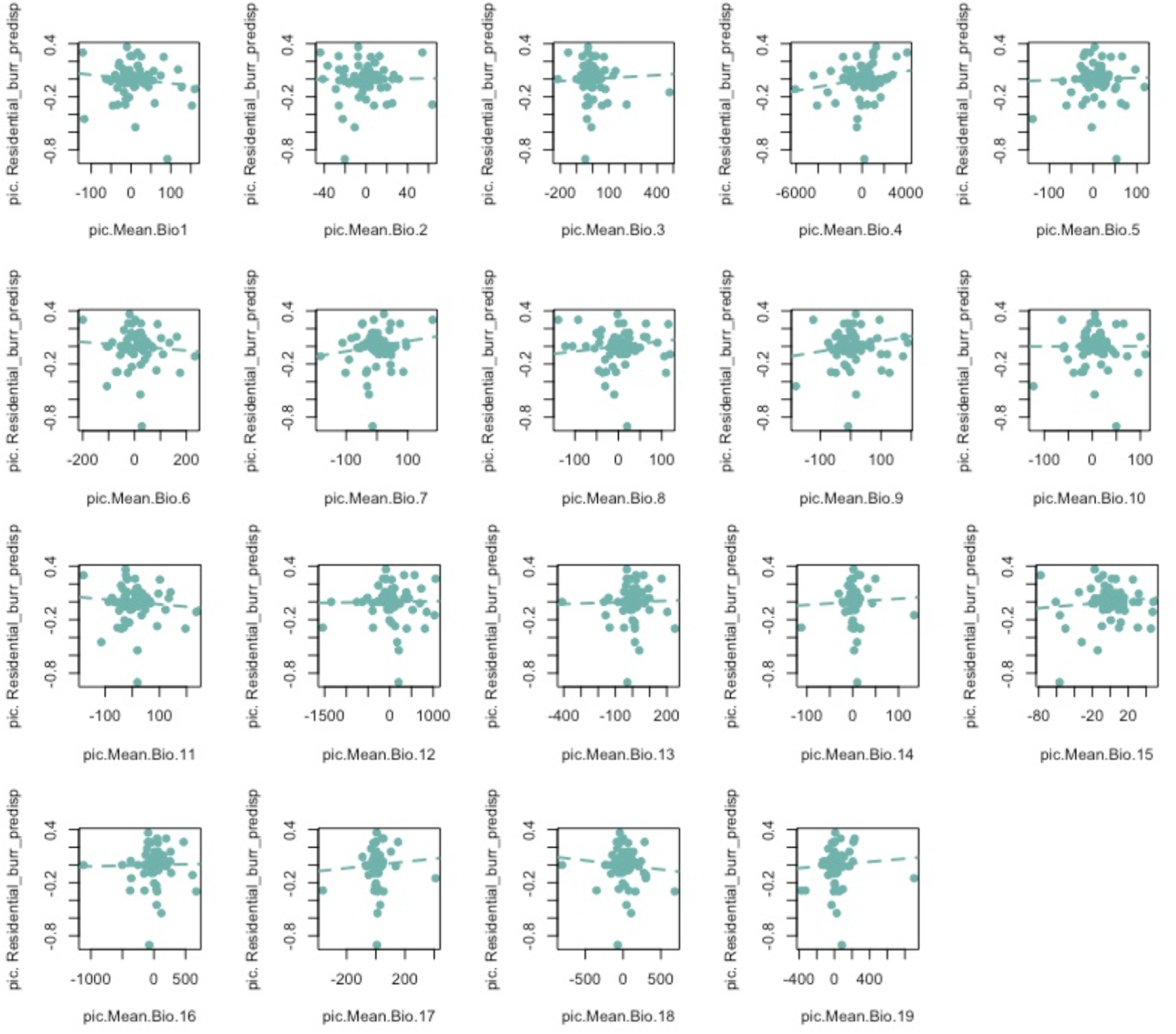
Phylogenetically controlled regressions of a residential burrowing predisposition against bioclimatic variables (number of offsprings/ litter: left top and left bottom; the number of litters: right top and right bottom).

#### Lifestyle and habitat openness (Hypothesis 4.1)

Habitat openness was not associated with burrowing behavior (Figure 9; burrows provide effective protection against predators, particularly in open habitats; PIC_habitat openness vs PIC_Burrowing propensity_Residential lifestyle - r-squared: -0.011, p-value > 0.05; PIC_habitat openness vs PIC_Burrowing propensity_Breeding lifestyle - r-squared: -0.018, p-value > 0.05). When four lifestyle states were considered, on average, different lifestyles were associated with similar habitats (Phylogenetic ANOVA, Residential lifestyle: F-value: 0.43 and p-value: 0.96; Breeding lifestyle: F-value: 0.13 and p-value: 0.99).

**Figure 9:**
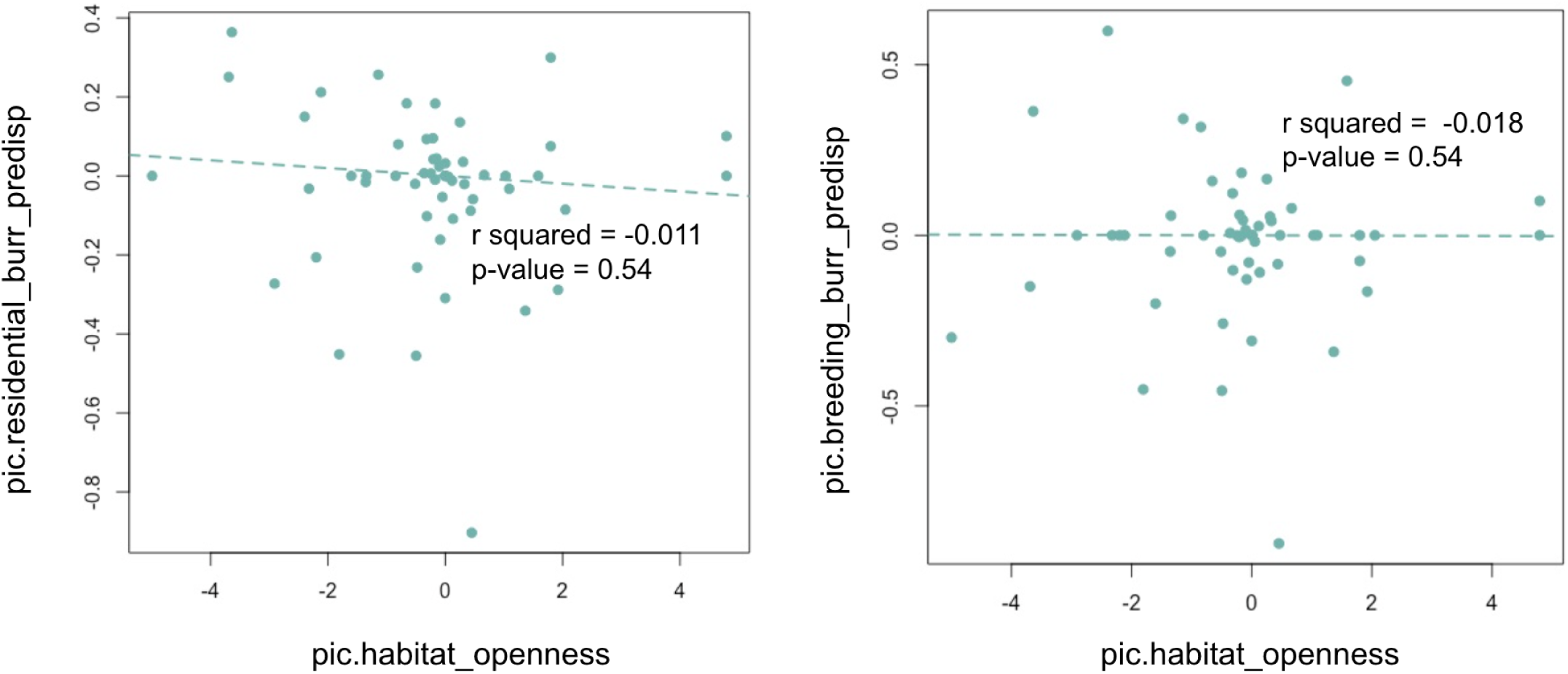
Phylogenetically controlled regressions of habitat openness against residential and breeding burrowing predisposition showing no relationship between variables

### Associations of life history with body size, climate, sociality, and lifestyle

#### Life history and allometric scaling (Hypothesis 5.1)

Small body-sized animals did not have higher fecundity (number of offspring/litter and number of litter) as expected (Figure 10)(Number of offspring/litter - r-squared: 0.007, p-value: 0.47; Number of litters - r-squared: 0.015, p-value: 0.78).

**Figure 10:**
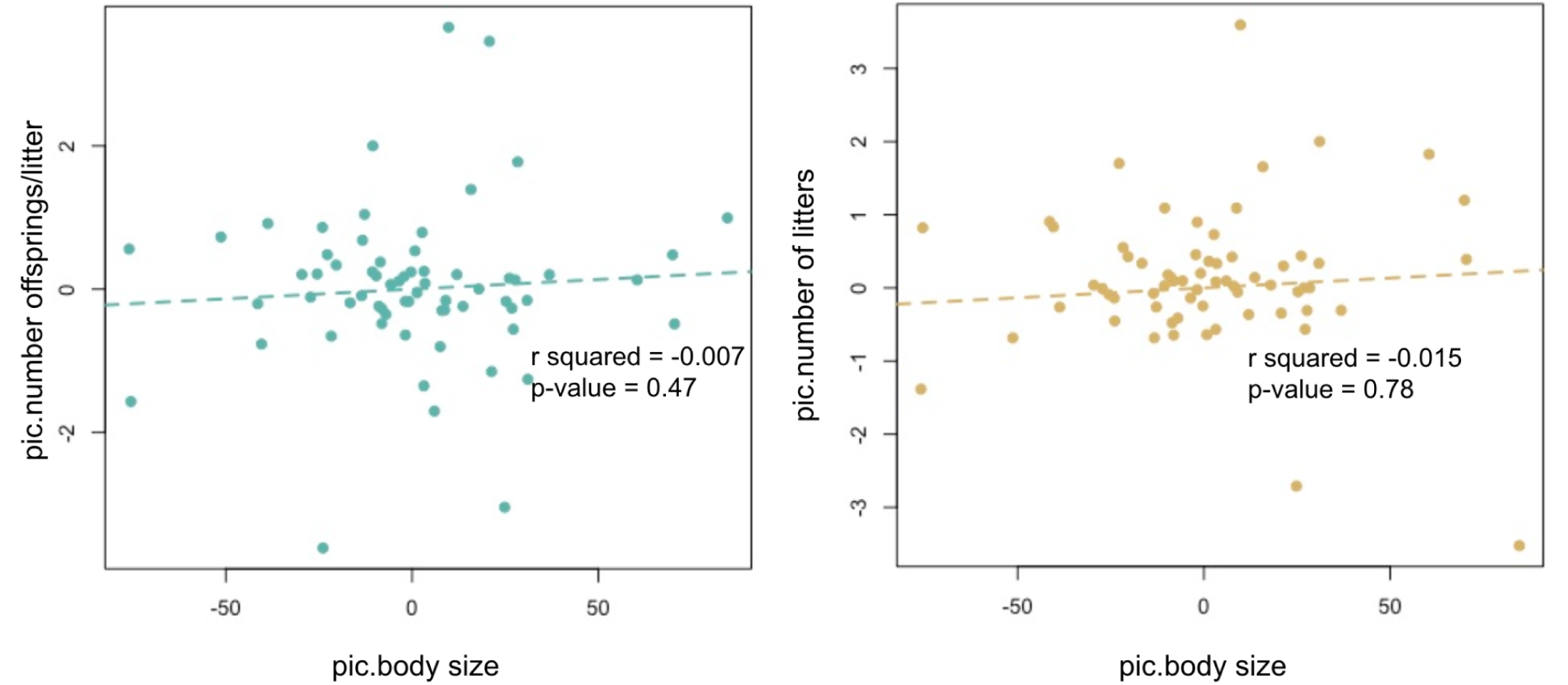
Phylogenetically controlled fecundity regressions against body size show no relationship between variables.

#### Life history and climatic regimes (Hypothesis 5.2)

The fecundity of species was weakly correlated with climate (Figure 11). As expected, seasonal descriptions of climatic envelopes (maximum temperature of the warmest month, mean temperature of warmest and driest quarter, precipitation of warmest quarter, temperature seasonality, annual temperature range, and mean temperature in the wettest quarter) were statistically significant in explaining fecundity traits. However, the effect sizes were pretty small (number of offspring/litter: 0.04-0.05, number of litters: 0.05-0.10) (Figure 11).

**Figure 11:**
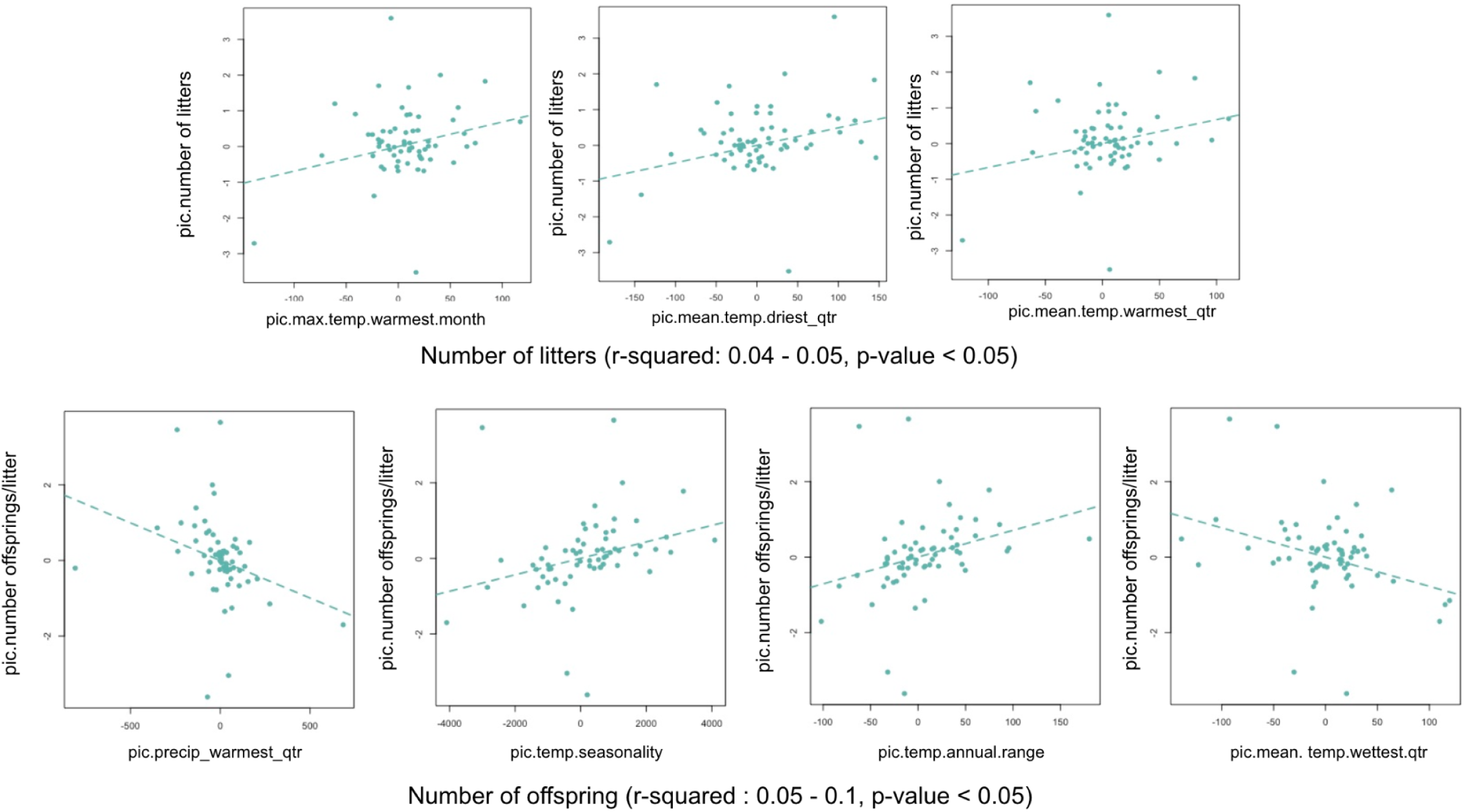
Phylogenetically controlled fecundity regressions against select climatic variables with low effect sizes.

#### Life history and sociality (Hypothesis 5.3)

Social species were more fecund than non-social species, where social species were associated with a larger number of offspring/litter (r-squared: 0.15, p-value: <0.05) but not with the number of litters (r-squared: 0.01, p-value: 0.58) (Figure 12).

**Figure 12:**
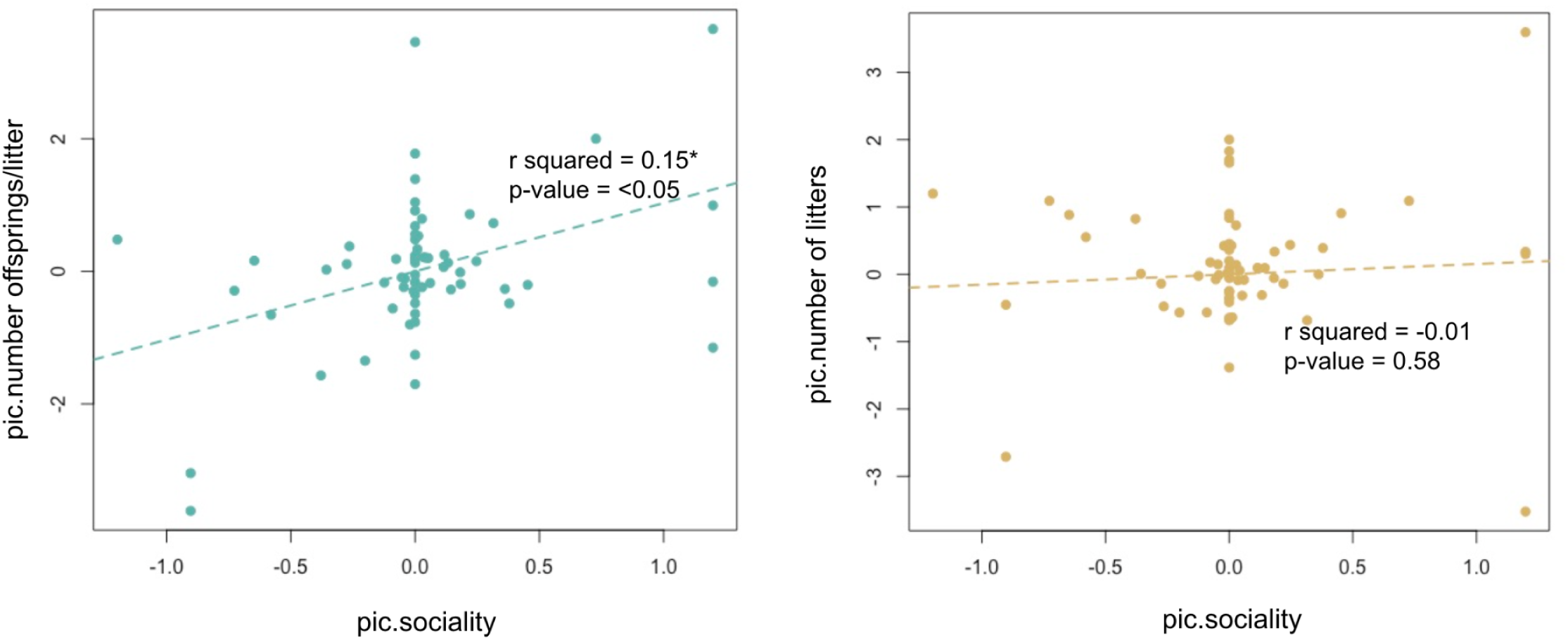
Phylogenetically controlled regressions of fecundity against sociality.

#### Life history and Lifestyle (Hypothesis 5.4)

Species of burrowing Lagomorphs also had a larger number of offspring/litter and a larger number of litters/year, suggesting an r-selected life history strategy (Figure 13). Additionally, the relationship between fecundity and burrowing was stronger with ‘number of offspring/litter’ than ‘number of litters’ (Figure 10; PIC_Burrowing propensity_Residential lifestyle vs. PIC_Number of offspring/litter- Slope: 3.17, r-squared: 0.3774, p-value < 0.05; PIC_Burrowing propensity_Residential lifestyle vs. PIC_Number of litters - Slope: 1.65, r-squared: 0.096, p-value < 0.05; PIC_Burrowing propensity_Breeding lifestyle vs. PIC_Number of offspring/litter- Slope: 2.4, r-squared: 0.2746, p-value < 0.05; PIC_Burrowing propensity_Breeding lifestyle vs. PIC_Number of litters - Slope: 0.69, r-squared: 0.007, p-value < 0.05). This seems to be largely accurate in Ochotonidae (and true rabbits), with burrowing species of pikas being more fecund and reaching early age maturity in comparison to rock-dwelling species (from the very few well-studied species of Lagomorphs we know life history patterns about) (Table S2_Final Data).

**Figure 13:**
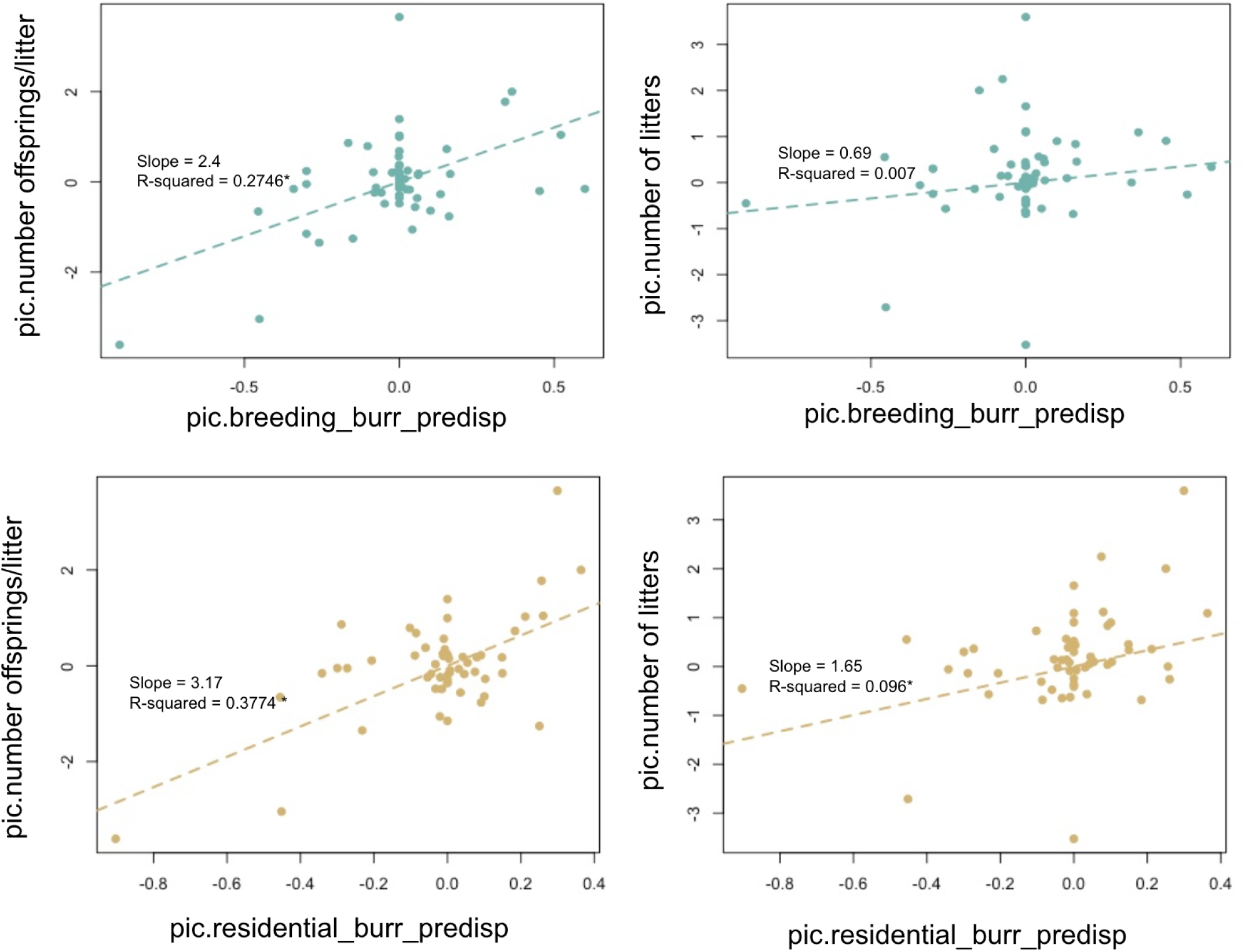
Phylogenetically controlled regressions of fecundity against burrowing

## Discussion

### Lagomorpha phylogeny

We constructed the most comprehensive, well-resolved, dated phylogeny of Lagomorpha sampling 84/102 taxa over and above most published studies (Cano-Sánchez et al. 2022; Kriegs et al. 2010; Ge et al. 2013). Since most recent studies failed to include all species of *Ochotonids*, we sequenced genomes of two Asian pikas (*Ochotona nubrica* and *Ochotona ladacensis*), the latter being the first genome for the species (Wang et al. 2020; Tang et al. 2022).

The placement of *Ochotona ladacensi*s has had a sketchy past with specimens in the past, being assigned to the neighboring black-lipped pika (*Ochotona curzoniae,* clade *Ochotona*) based on morphological similarity, although most authors now consider it as a separate entity (Smith et al. 2010; Menon 2014; Smith et al. 2018) While being morphologically closest to the Pallas pika (*Ochotona pallasi*, clade *Pika*), it differs from it in skull morphology (tympanum bullae and cranium curvature) (Chapman et al. 1990). It was first placed in the clade *Conothoa* based on mitochondrial cytb sequences, although its relative placement within the clade needed to be better resolved (Niu et al. 2004). Subsequently, its placement in the clade *Conothoa* was confirmed and inferred to be a part of the O.roylei and O.macrotis group based on more cytb sequencing (Lissovsky 2014). A preliminary examination of skull morphology indicated that the species had unique morphology and was distinct from other species (only one skull of *O.koslowi* lay within the canonical space of *O. ladacensis*) (Lissovsky 2014). A follow-up investigation of skulls from animals of the clade *Conothoa,* the phylogenies of both mitochondrial (cytb + COI) and nuclear markers (ILRAPL1, MGF, OXA1L, TTN, RAG1, RAG2) found weak to-moderate support to place *O.ladacensis* in the western pika In sub-clade (Himalayas, Pamir-Tian Shan mountains) along with *O.macrotis, O.roylei, O.rutila,* and *O.rufescens* (Lissovsky et al. 2022). We resolved the placement of the species and infer its position in the clade *Conothoa* basal to the *Ochotona himalayana, Ochotona macrotis,* and *Ochotona roylei* group.

We examined the evolution of burrowing behavior, its adaptive significance, and its associations with other behavioral traits using a combination of ancestral state reconstructions and the comparative method.

### Ancestral Reconstruction of Lifestyle and Morphometric Traits

Ancestral state reconstructions of lifestyle revealed that the ancestor of all *Ochotonids* was a burrowing species rather than a rock-dwelling species, as suggested by other research (Feijó et al. 2020). This is probably because our study sampled a clade of burrowing animals (*Alienouroa)* that was recently sequenced and was missed by this earlier study (Feijó et al. 2020; Wang et al. 2020). In addition, our categorization of lifestyles was done systematically and at several different levels to ensure that our findings were not an artifact of our classification of traits (see methods). The ancestor of all *Leporids* was inferred to be either living in forms/burrows of other species. Subsequently, there have been several transitions to rock-dwelling behavior in *Ochotonidae (*subclade *Ochotona* and *Conothoa)* and once in *Leporidae (Pronolagus* hares). There have also been a few instances of transitions to burrowing behavior from forms/burrows of other species in *Leporidae* and many transitions to flexible lifestyles.

Ancestral reconstructions of body size revealed that the ancestor of all Lagomorphs had an intermediate body size and that there has been one instance of small body-sized evolution in *Ochotonidae* and one instance of large body-size evolution in *Leporidae* (true-hares) as suggested earlier (Ge et al. 2013). Body size evolution over and above genetic drift is also likely to have been shaped by competition and predation in open habitats (Tomiya and Miller 2021).

### Examining correlations with lifestyle

In this study, we find that burrowing was moderately associated with sociality. This lends support to the hypothesis that burrowing is energetically expensive, with sociality offsetting the costs of burrowing as seen in many systems (Hypothesis 2) (Spinks 1998; Ebensperger and Bozinovic 2000; Ebensperger and Cofré 2001; Lacey and Wieczorek 2003; Walker et al. 2007; McAlpin et al. 2011). However, we did not find evidence for other hypotheses, such as - morphological adaptations for scratch-digging in burrowing species (Hypothesis 1) (Emlen and Keith Philips 2006; Macagno et al. 2016; Gomes Rodrigues and Damette 2023; Rodrigues et al. 2023), burrowing as behavioral adaptations to extreme/variable climate (Hypothesis 3.1, 3.2) (Nikol’skii and Khutorskoi 2001; Belovezhets and Nikol’skii 2012; Pike and Mitchell 2013; Milling et al. 2018), and burrowing as behavioral adaptations to predation pressure in open habitats (Hypothesis 4) (Lima and Dill 1990; Bonenfant and Kramer 1996; Harper and Batzli 1996; Jackson 2000; Hemmi 2005a; Hemmi 2005b; Wilson et al. 2012) although studies with other model systems have found such patterns.

### Examining correlations with life history

The fecundity of Lagomorphs was moderately associated with sociality, with social species having a larger number of offspring/litter but not a larger number of litters. This supports the hypothesis that social species are generally more fecund, as documented earlier (Hypothesis 5.3) (Ebensperger et al. 2012). Burrowing species were also more fecund, probably stemming from stable thermal environments (Pike and Mitchell 2013; Millar et al. 2016; Milling et al. 2018) or better protection against predators (Lima and Dill 1990; Hemmi 2005; Hemmi 2005; Wilson et al. 2012) that leaves behind a larger fraction of the energy acquired to be allocated for reproduction (Hypothesis 5.4) (Alves et al. 2007). Fecundity patterns were weakly associated with extreme/seasonal bioclimatic variables as expected where climate constrained the (Hypothesis 5.2) (Hufnagl et al. 2011; Battistella et al. 2018; Verhagen et al. 2020). Interestingly, allometric scalings were not pronounced in Lagmorphs, and body size poorly explained fecundity patterns (Hypotheses 5.1) (Pianka 1970; Blueweiss et al. 1978; Sibly and Brown 2007). Our findings indicate how burrowing, sociality, and fecundity are associated with each other in Lagomorphs (Figure 14; Burrowing-Sociality-Fecundity triangle). It is likely that such patterns also hold true for other taxa, such as rodents, where burrowing species are social and r-selected when compared to species with other lifestyles.

**Figure 14:**
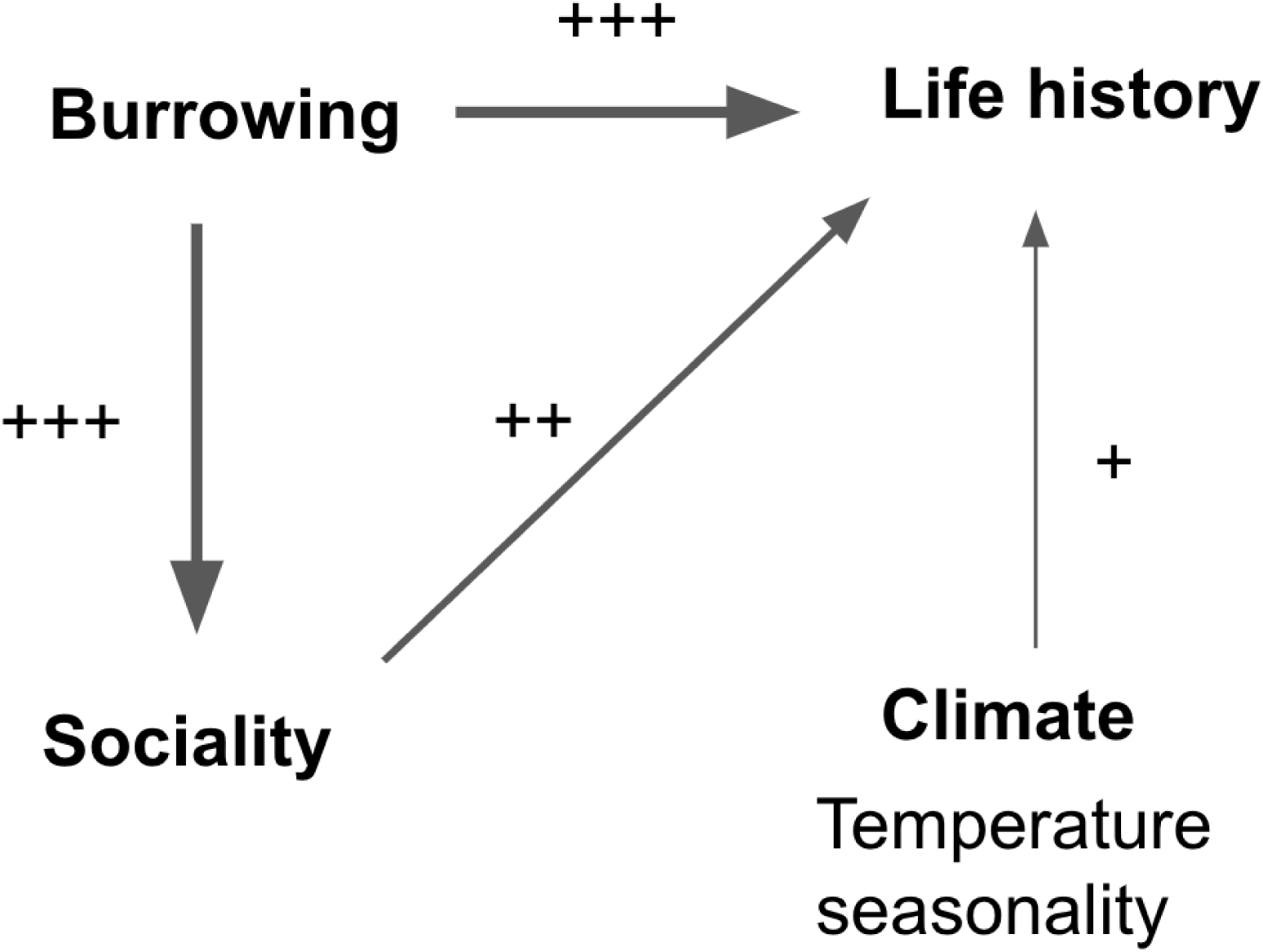
Burrowing-Sociality-Fecundity triangle describing associations between traits. The thickness of the arrows and the number of ‘+’ marks indicate the strength of relationships, as seen in Lagomorpha.

We constructed the first comprehensive phylogeny of Lagomorpha and examined the evolution of lifestyle, sociality, life history, and their adaptive potential using a combination of ancestral state reconstructions and the comparative method. In the process, we built a large database collating data from various sources, which is a large feat, accounting for six long months of work. We found complex patterns of burrowing evolution with transitions to burrowing from other states and to other states from burrowing. While we recovered patterns between burrowing and sociality, sociality, and fecundity known from other studies in rodents, we also recovered the burrowing-sociality-fecundity triangle. This highlights the need for large-scale comparative phylogenetic studies that examine adaptations on a large time scale (historic timescales) in conjunction with experimental studies over small time scales (current timescales).

## Supporting information

Supplementary Fgures

## Acknowledgments

We thank Dr. Yash Veer Bhatnagar, Dr. Nagarjun Vijay, and Dr. Robin Vijayan for their valuable comments and suggestions on the manuscript. We are also grateful to members of the Sciurid Lab and the Bird Lab at IISER Tirupati for their feedback on the manuscript, particularly Dr. Bishwarup Paul, Senan D’Souza, Dr. Swathi Udayraj, Dr. Nivetha Murugesan, Chiti Aravind, Dr. Naman Goyal, Dr. Meghana Natesh, and Vinay K. L. for numerous discussions on various aspects of this work.

