## Supplementary Fgures for "Evolution of burrowing and associated behavioral traits in Lagomorphs"

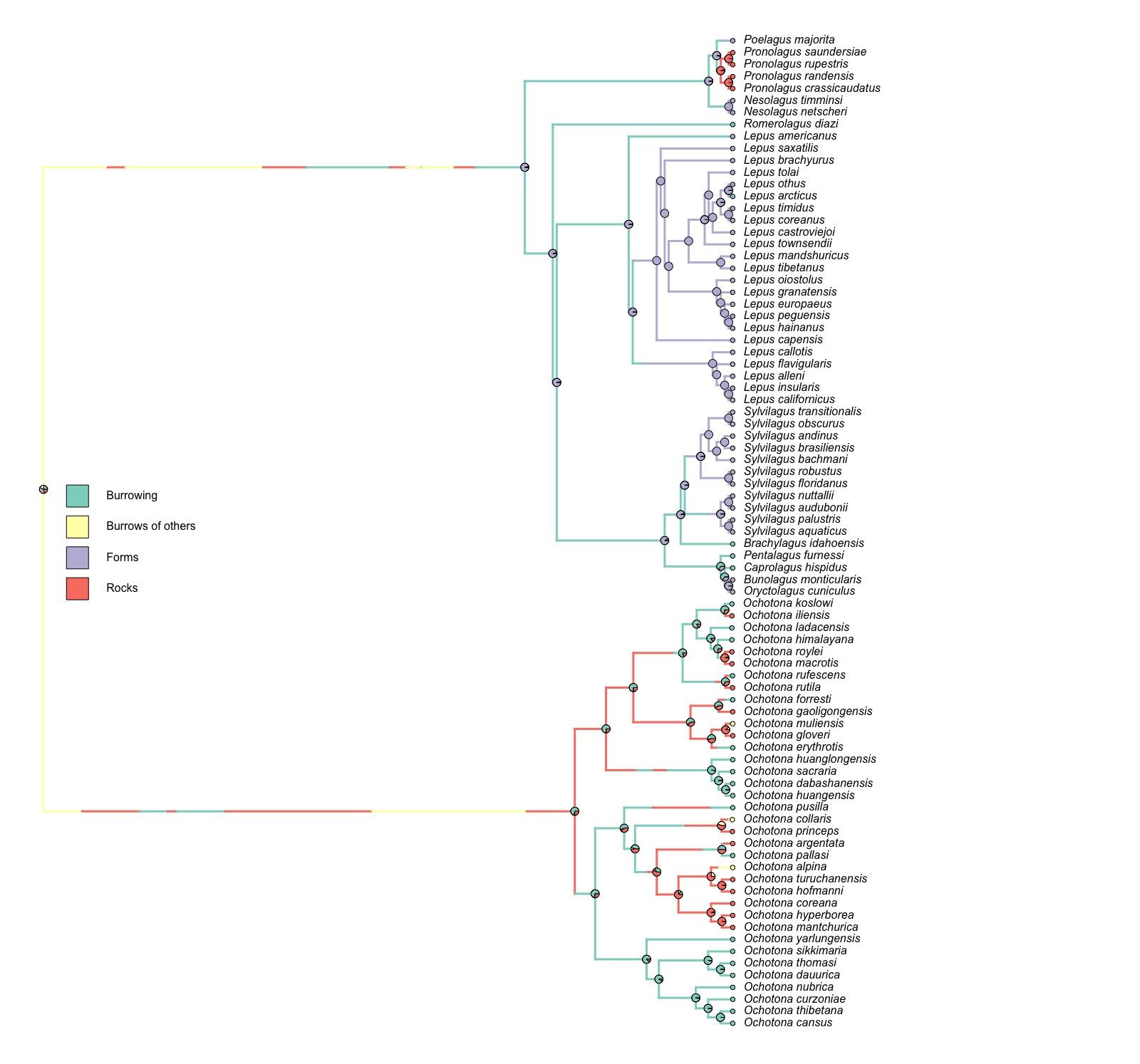

Figure S1: Ancestral state reconstructions of residential lifestyle with four trait states. The most energetically expensive state (Burrowing > Burrows of others > Forms > Rocks) was used to represent the residential lifestyle state of species.

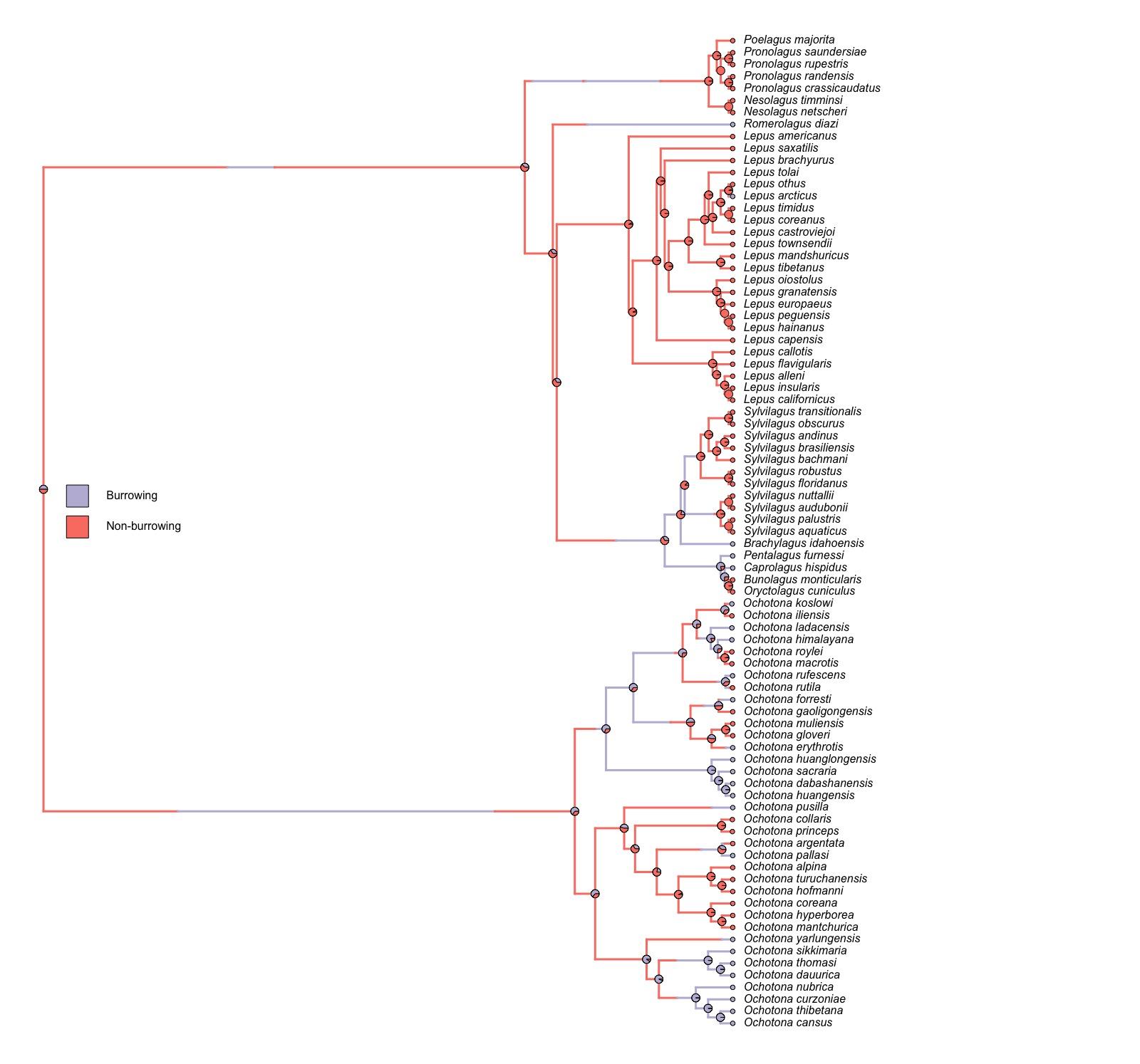

Figure S2: Ancestral state reconstructions of residential lifestyle with two trait states. Only two trait states (burrowing and non-burrowing) were used to represent the residential lifestyle state of the species.

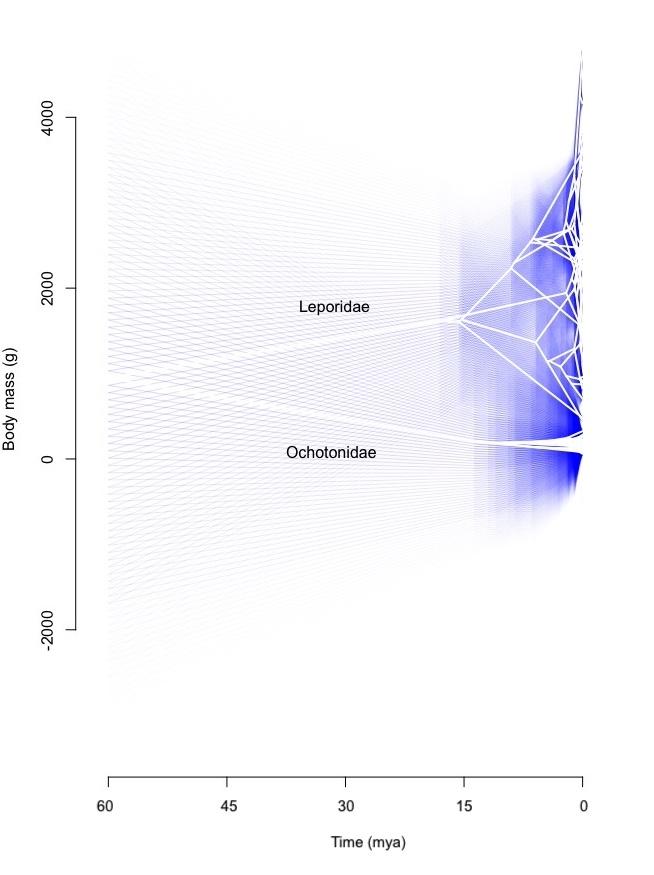

Figure S3: Phenogram of body mass changes along phylogeny with 95% confidence intervals overlaid.

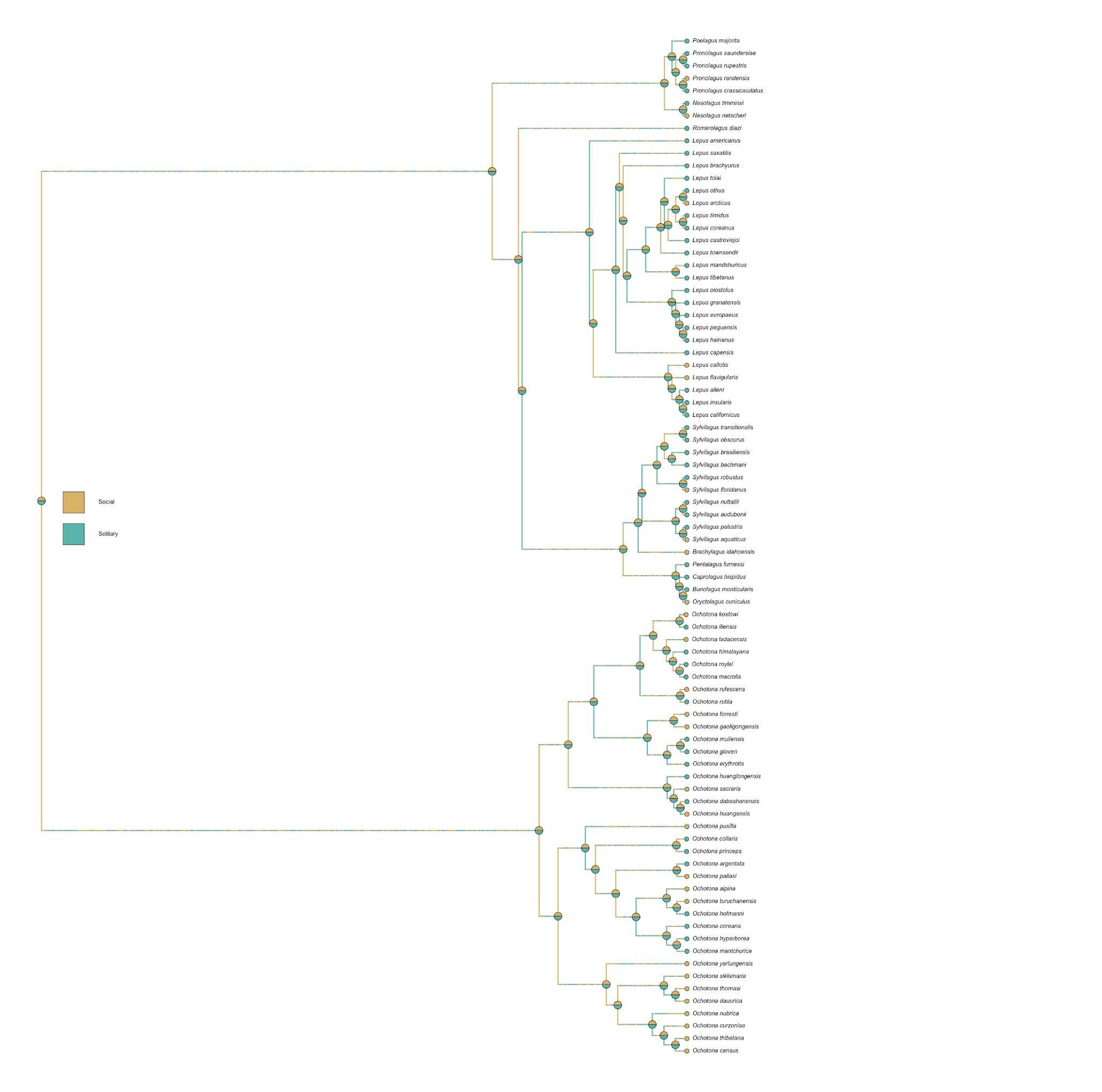

Figure S4: Reconstructions of sociality showing poor resolution of ancestral states.
